# Consequences of single-locus and tightly linked genomic architectures for evolutionary responses to environmental change

**DOI:** 10.1101/2020.01.31.928770

**Authors:** Rebekah A. Oomen, Anna Kuparinen, Jeffrey A. Hutchings

**Affiliations:** Centre for Ecological and Evolutionary Synthesis, University of Oslo, Oslo, Norway; Centre for Coastal Research, University of Agder, Kristiansand, Norway; Department of Biological and Environmental Sciences, University of Jyväskylä, Jyväskylä, Finland; Department of Biology, Dalhousie University, Halifax, NS, Canada; Institute of Marine Research, Flødevigen Marine Research Station, His, Norway

**Keywords:** climate change, evolutionary simulation, genetic architecture, linkage disequilibrium, recombination rate, structural genomic variation

## Abstract

Genetic and genomic architectures of traits under selection are key factors influencing evolutionary responses. Yet, knowledge of their impacts has been limited by a widespread assumption that most traits are controlled by unlinked polygenic architectures. Recent advances in genome sequencing and eco-evolutionary modelling are unlocking the potential for integrating genomic information into predictions of population responses to environmental change. Using eco-evolutionary simulations, we demonstrate that hypothetical single-locus control of a life history trait produces highly variable and unpredictable harvesting-induced evolution relative to the classically applied multi-locus model. Single-locus control of complex traits is thought to be uncommon, yet blocks of linked genes, such as those associated with some types of structural genomic variation, have emerged as taxonomically widespread phenomena. Inheritance of linked architectures resembles that of single loci, thus enabling single-locus-like modeling of polygenic adaptation. Yet, the number of loci, their effect sizes, and the degree of linkage among them all occur along a continuum. We review how linked architectures are often associated, directly or indirectly, with traits expected to be under selection from anthropogenic stressors and are likely to play a large role in adaptation to environmental disturbance. We suggest using single-locus models to explore evolutionary extremes and uncertainties when the trait architecture is unknown, refining parameters as genomic information becomes available, and explicitly incorporating linkage among loci when possible. By overestimating the complexity (e.g., number of independent loci) of the genomic architecture of traits under selection, we risk underestimating the complexity (e.g., nonlinearity) of their evolutionary dynamics.

## Eco-evolutionary responses hinge on genetic and genomic architecture

Predicting the responses of populations and species to anthropogenic disturbance is a major challenge, and one that requires urgent attention given the current climate and biodiversity crises (IPCC 2018; IPBES 2019). Advances in second- (high throughput) and third- (long read) generation sequencing have produced a wealth of sequence and structural genomic data on non-model organisms. These data have great potential to inform eco-evolutionary models of the responses of natural populations to a variety of selection pressures (Hoffmann *et al.* 2015; Bay *et al.* 2017a; Coulson *et al.* 2017). Such models can inform current management strategies and facilitate planning for future environmental conditions and associated ecosystem structures.

Key parameters influencing evolutionary responses are the genetic/genomic architectures underlying adaptive traits. These refer to how a trait is controlled by one or more genes and interactions among alleles (e.g., number and effect sizes of contributing loci, dominance, epistasis, pleiotropy), structural arrangement (e.g., inversions, fusions, translocations, duplications), position, and linkage among loci. These characteristics contribute to the inheritance models for genes underlying adaptive traits that are used in evolutionary predictions.

Here, we discuss some effects of genetic (the number of loci and their effect sizes) and genomic (the degree of linkage among loci) architectures on predictions of evolutionary responses to environmental disturbance. Single or unlinked loci have received comparatively more attention than linked genomic architectures in this regard (Bay *et al.* 2017a; Kardos & Luikart 2019). We demonstrate that hypothetical single locus control of a life history trait under harvesting-induced selection generates a more variable evolutionary response compared to an unlinked polygenic scenario. We then suggest that linked polygenic architectures resemble those of single large-effect loci, yet exist along a continuum of linkage disequilibrium, and are likely to play a large role in adaptation to rapid environmental change. We show that linked architectures underlie diverse traits in natural populations that are directly or indirectly under environmental selection. Finally, we discuss some barriers to modelling such architectures and challenges they present to conservation and management. More broadly, we aim to promote the integration of genomic data into eco-evolutionary modelling of responses to environmental change.

### Large-effect loci alter evolutionary predictions compared to traditional polygenic models

The degree to which a trait is controlled primarily by a single locus or multiple loci will influence its evolution in response to environmental stressors. Traditional evolutionary models have focused on the fixation dynamics of single-locus traits suddenly exposed to selection (Orr & Unckless 2014). As single-locus control of complex traits has been considered rare (Feder & Walser 2005), eco-evolutionary models of complex non-model organisms often incorporate a standard inheritance model of 10 or 20 unlinked loci (e.g., Kuparinen & Hutchings 2012). Multi-locus (e.g., 100+ loci) models based on genomic single nucleotide polymorphism (SNP) data have also been employed more recently to predict the capacity of a population to evolve in pace with global climate change (Bay *et al.* 2017a, 2018). Bay *et al.* (2017b) described a potential framework for genomic predictions of adaptive responses to environmental change, termed ‘evolutionary response architectures’, but focused on unlinked polygenic control of climate-associated traits.

The assumption that single genetic variants accounting for large amounts of phenotypic variation are rare is being challenged as more refined statistical genomics enable their discovery (Hoban *et al.* 2016). Large-effect loci have been documented in plants (Kivimäki *et al.* 2007; Baxter *et al.* 2010), insects (Reed *et al.* 2011), mammals (Johnston *et al.* 2013; Kardos *et al.* 2015; Jones *et al.* 2018; Barrett *et al.* 2019), birds (Toews *et al.* 2019; Merritt *et al.* 2020), and fishes (Colosimo *et al.* 2004; Lampert *et al.* 2010; Barson *et al.* 2015; Thompson *et al.* 2019). For example, the *optix* gene controls wing pattern in *Heliconius* butterflies (Reed *et al.* 2011), the *RXFP2* gene controls horn architecture in Soay sheep (Johnston *et al.* 2013), and the *Agouti* locus controls coat colour polymorphism in deer mice (Barrett *et al.* 2019). The implications of these variants for eco-evolutionary model predictions can be severe. After the discovery that the *vgll3* gene is responsible for 40% of the variation in age at maturity in Atlantic salmon (*Salmo salar*) (Barson *et al.* 2015), Kuparinen and Hutchings (2017) demonstrated that hypothetical single-locus control of this key, sexually dimorphic life-history trait generates divergent and disruptive fisheries-induced evolution relative to that predicted by the classically applied, commonly assumed multi-locus model. As demonstrated in Box 1, these chaotic dynamics are largely driven by the single-locus control, not the sexual dimorphism, which indicates that they could occur for a broad range of traits. This finding is consistent with recent simulations demonstrating increased variability in population viability under rapid directional environmental change when the trait under selection is under single-locus rather than multi-locus control (Kardos & Luikart 2019). It further suggests that recovery following relaxation of selection is also highly variable.

### Inheritance of linked polygenic architectures resembles that of single loci

Particularly for complex traits, such as those contributing to growth, behaviour, or environmental responses, polygenic control might indeed be the norm (Savolainen *et al.* 2013; Palumbi *et al.* 2014; Bay *et al.* 2017a). Yet, blocks of tightly linked putatively adaptive genes that undergo reduced or no recombination are taxonomically widespread (Nosil *et al.* 2009; Rogers *et al.* 2011; Yeaman 2013; Küpper *et al. 2016;* Wellenreuther & Bernatchez 2018; Pearse *et al.* 2019).

There are several mechanisms by which linked clusters might evolve in response to selection, but all are characterized by a reduction in recombination (Yeaman 2013). This is because when linkage captures an advantageous allelic combination, selection will favour a reduced recombination rate in that region to avoid splitting up complementary alleles (Charlesworth & Charlesworth 1979; Kirkpatrick & Barton 2006; Bürger & Akerman 2011; Yeaman & Whitlock 2011). Low recombination rates can be achieved by genic modifiers that decrease the frequency of crossovers during meiosis or genomic rearrangements that alter gene order, suppress recombination, and/or generate unbalanced gametes in heterozygotes (Figure 1; Butlin 2005; Ortiz-Barrientos *et al.* 2016). Recombination rate is also negatively correlated with epistasis, chromosome length, and proximity among loci (Figure 1; Kong *et al.* 2002; Butlin 2005). Sequence content can have both positive (CpG content) and negative (GC, polyA/polyT, and heterochomatin content) effects on recombination rate (Figure 1; Kong *et al.* 2002).

**Figure 1:**
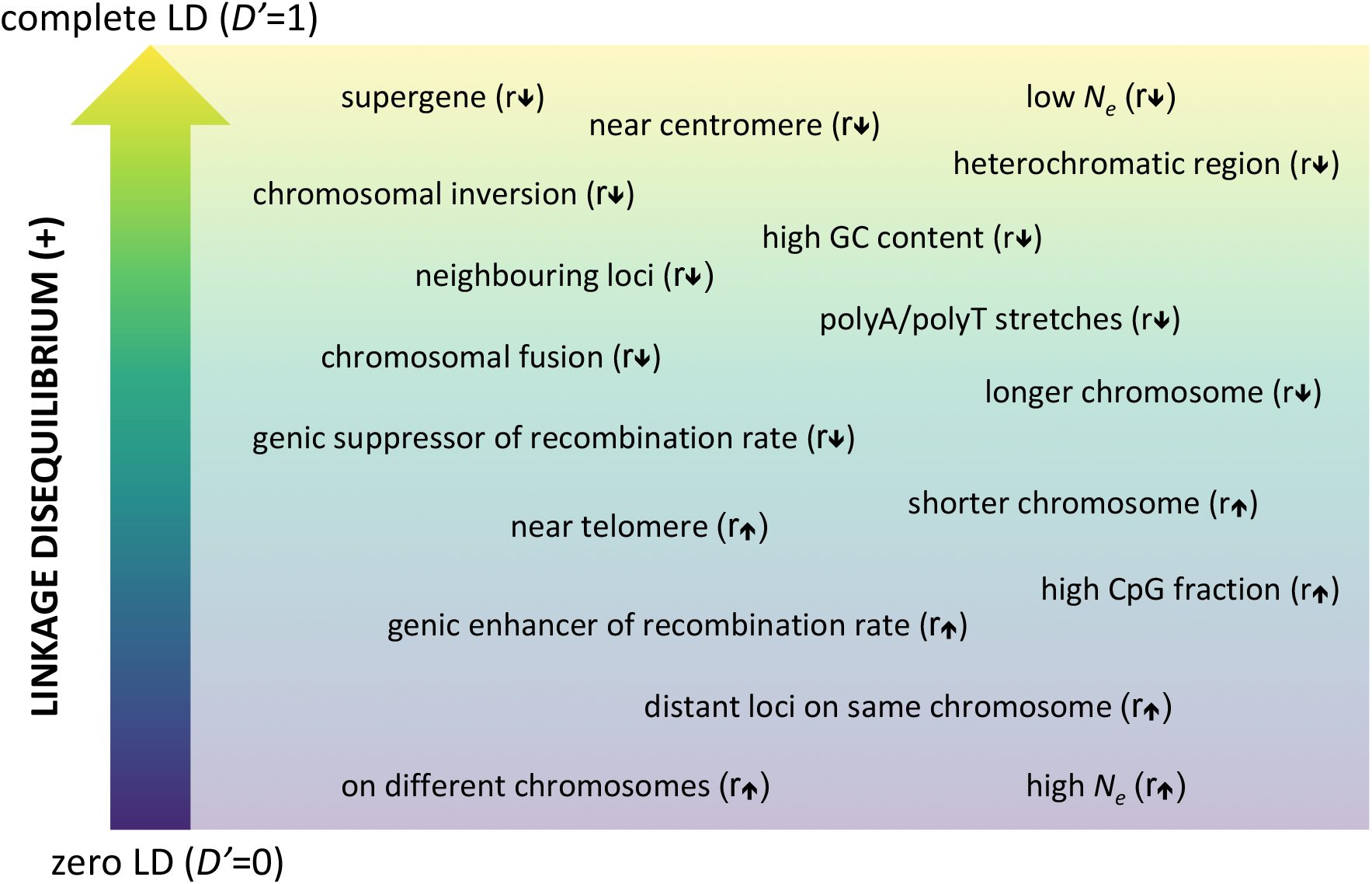
Positive linkage disequilibrium (LD) exists on a spectrum influenced by several factors potentially affecting recombination rate (r) among loci. *D’* is a normalized metric of LD. For positive LD it is represented by the function *D’* = (x_aa_ - p_a_ × q_a_)/min(x_ab_, x_ba_), whereby x is haplotype frequency and p and q are allele frequencies for two polymorphic loci with two alleles (a, b) each. Rather than a mechanistically accurate diagram, consider this figure as a roughly organized corkboard onto which the various factors and conditions have been pinned.

The lower the rate of recombination, the greater the degree of linkage disequilibrium (LD; non-random association of alleles at different loci). LD is also inversely proportional to effective population size through its relationship to genetic drift (Sved 1971; Waples *et al.* 2016), which could have notable consequences for small populations, as they would tend to harbour loci in higher LD. When LD is high, the inheritance pattern of a genomic region containing multiple loci resembles that of a single locus, such that complete linkage among genes would result in their co-inheritance (Figure 2). While, historically, precise estimates of LD have been challenging to obtain and are therefore not readily available for most species or populations, accessible high-throughput sequencing of non-model organisms is rapidly eliminating this barrier (Box 2). Other barriers to integrating linkage into eco-evolutionary models include a lack of understanding of the potential fitness effects of regions of high LD (e.g., partial sterility of heterozygotes, accumulation of deleterious mutations; Box 2).

**Figure 2:**
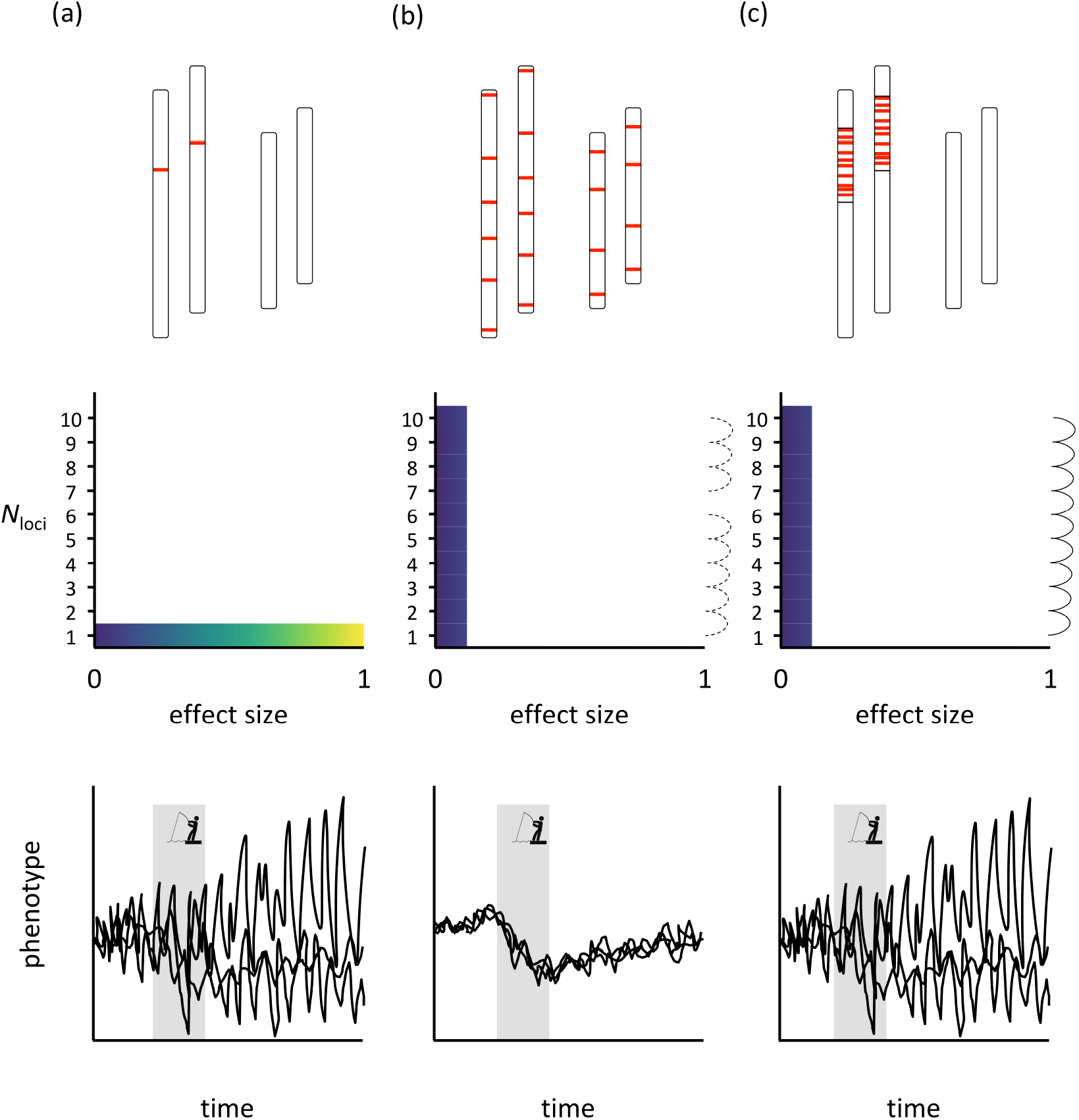
Hypothetical genomic architectures (top row), corresponding model parameters (middle row), and evolutionary simulations (bottom row) for a polygenic trait in a diploid individual with two chromosomes that is under a temporary period of directional selection (grey bars): (a) a single locus of large effect, (b) ten loci of small effect with negligible linkage disequilibrium (LD), and (c) ten loci of small effect with strong LD. Red bars indicate the position of individual loci along a chromosome. Model parameters include the number of loci (*N*_loci_), their effect sizes, and the degree of LD among them (when applicable), whereby continuous dashed lines indicate negligible levels of LD and continuous solid lines indicate strong LD. Individual black lines represent hypothetical replicate simulations (*N*=3).

### Linked polygenic architectures are ubiquitous and exist along a continuum of linkage disequilibrium

Linked regions are often identified in population genomic studies as ‘genomic islands of divergence’ (regions that exhibit greater differentiation than expected under neutrality; Wu 2001), although the degree of linkage varies depending on the mechanism of recombination suppression (Figure 1). For example, genic modifiers might only partially reduce recombination, leading to low or moderate levels of LD (Butlin 2005).

In contrast, extreme cases of tightly linked co-adapted gene complexes associated with discrete complex phenotypes, known as supergenes, underlie key life-history traits in a variety of species (Schwander *et al.* 2014). Supergenes are often associated with structural genomic variation, which underlies complex phenotypes and adaptive processes in a wide variety of non-model taxa (Mérot *et al.* 2020; Wellenreuther *et al.* 2019). Non-recombining sex chromosomes are extreme examples of supergenes extending entire chromosomes, many of which evolved through structural genomic mutations (e.g., the mammalian Y chromosome evolved through a series of inversions following the formation of the Sex-Determining Region Y gene) (Lahn & Page 1999; Bachtrog 2013). There is emerging evidence that structural genomic variants might comprise the most important source of genomic variation in natural populations (Mérot *et al.* 2020), as they have been found to account for several times more variation, in terms of the number of affected nucleotides, than SNPs (e.g., 3× in the Australasian snapper [*Chrysophrys auratus*; Catanach *et al.* 2019], 12× in *Homo sapiens* [Pang *et al.* 2010]).

One of the most well studied types of structural variant is the chromosomal inversion (Sturtevant 1921; Wellenreuther *et al.* 2019). Inversions prevent recombination within the inverted region by displacing crossovers away from breakpoints during meiosis (at least in *Diptera* flies) or by producing lethal meiotic products (if the inversion does not include the centromere) or inviable gametes (if the inversion spans the centromere) in heterokaryotypes, resulting in the selective recovery of non-recombinant chromosomes (Reiseberg 2001; Hoffmann & Rieseberg 2008; Wellenreuther & Bernatchez 2018).

Other types of structural variation, such as chromosomal fusions and fissions, translocations, and copy number variants (CNVs), can also generate unbalanced gametes (Rieseberg 2001), which is expected to have similar implications for recombination rate reduction. Chromosomal fusions reduce recombination to a lesser extent than inversions, but to varying degrees in both heterozygotes and fused homozygotes (Bidau *et al.* 2001; Guerrero & Kirkpatrick 2014). There is also at least one example of a complex CNV maintaining linkage among candidate genes associated with multiple traits, effectively acting as a supergene (Tigano *et al.* 2018). While the precise mechanisms for reducing recombination are not yet clear for many types of structural variation, the field is poised for major advances (Mérot *et al.* 2020).

### Linked architectures play a large role in adaptation to rapid change

Linked architectures are hypothesized to facilitate rapid adaptation by enabling inheritance of co-adapted gene complexes. Instead of accumulating beneficial alleles over multiple generations, they come as a package that has the potential to spread rapidly through a population, similar to a single large-effect gene (Kirkpatrick & Barrett 2015). Therefore, while the extent of recombination in linkage blocks can vary, a linked genomic architecture would enable single-locus-like modeling of polygenic adaptation. Their prevalence, especially the rising ubiquity of structural variation, necessitates a reassessment of common assumptions regarding the degree to which genes contributing to a polygenic trait are likely to be physically linked and/or experience reduced recombination.

### ‘Mixed-effect’ architectures only partially alleviate uncertainty

There are numerous examples of single loci (e.g., Carter 1977; Daetwyler *et al.* 2014; Carlson *et al.* 2016; Barrett *et al.* 2019) and blocks of tightly linked loci (e.g., supergenes; Schwander *et al.* 2014) controlling alternative phenotypes in a seemingly discrete, ‘monogenic’ manner. Yet, mixed architectures consisting of a large-effect locus, supergene, or haploblock in addition to numerous small-effect, potentially unlinked, loci are likely common as well (herein, referred to as ‘mixed-effect architectures’). Small-effect variants accompanying those of large effect are challenging to detect using common approaches (e.g., genome-wide and gene-environment associations) given the difficulties of distinguishing weak signatures of selection from demographic processes or selection on other traits (Hoban *et al.* 2016; Stephan 2016). Thus, whether large-effect variants are truly monolithic is especially difficult to confirm outside of domesticated and model species owing to diverse genomic backgrounds and environmental influences.

Sinclair-Waters *et al.* (2020) recently characterized the mixed-effect architecture of age-at-maturity in Atlantic salmon, using an extensive genome-wide association study of 11,166 males from a single aquaculture strain, combined with high-density SNP arrays and pedigree information. Including the previously known large-effect *vgll3* and *six6* loci, they identified 120 genes contributing to age-at-maturity with various effect sizes. One would expect such mixed-effect architectures to exhibit evolutionary dynamics intermediate to those of single locus and highly polygenic scenarios: the addition of many small-effect loci along with a large-effect locus should reduce the stochasticity and trait variance within and between populations. Indeed, Kardos & Luikart (2019) simulated a range of mixed-effect architectures underlying a phenotypic response to a sudden environmental shift and obtained intermediate levels of average phenotype, population viability, and extinction rate relative to the single locus and highly polygenic models. Therefore, while mixed-effect architectures seem to generate somewhat more predictable evolutionary dynamics compared to single locus architectures, they are unlikely to completely alleviate the concerning degree of stochasticity that seems to be characteristic of architectures with major effect loci. Therefore, the evolution of age-at-maturity in Atlantic salmon is still likely to exhibit increased variability relative to the classical multi-locus model of genomic architecture, but unlikely to be as extremely varied as the hypothetical single-locus scenario presented previously (Kuparinen & Hutchings 2017).

Unfortunately, such large-scale, high-throughput approaches as used for Atlantic salmon (Sinclair-Waters *et al.* 2020) are not feasible for most non-model species, which typically lack pedigree information and sufficient sample sizes and genomic resources, especially when they are of conservation concern. When the precise architecture is unknown, the genomic background (e.g., population or ecotype) in which a large-effect variant occurs can be considered as to whether it potentially influences expression of the focal variant for the trait of interest, but only if the variant is polymorphic within different genomic backgrounds. If the genomic background alters trait expression within particular genotypes of the focal variant (and independently of environmental factors), then there must be additional loci affecting the trait. How best to model mixed-effect architectures when they can be characterized requires further investigation and we encourage research in this direction. Nonetheless, in the next section, we seek to highlight the broad potential of modelling tightly linked architectures as single loci for the purpose of predicting responses of natural populations to environmental disturbance. Acknowledging the prevalence and ubiquity of large-effect linked architectures, in addition to single loci of large effect and mixed-effect architectures, is a critical next step towards modelling the full spectrum of genomic architectural complexity.

## Linked genomic architectures underlie diverse traits in natural populations that are directly or indirectly under environmental selection

In recent years, linked genomic architectures have been associated with a variety of adaptive traits in natural populations (Table 1). While inversions appear to be the most commonly studied (Wellenreuther & Bernatchez 2018; Wellenreuther *et al.* 2019), chromosomal fusions (Wellband *et al.* 2019) and complex architectures involving multiple rearrangements (Tigano *et al.* 2018; Pearse *et al.* 2019) are also associated, directly or indirectly, with traits relevant for adaptation. In some cases, SNPs located in proximity to one another are found to be in LD and the structural architecture is yet to be determined (e.g., Micheletti *et al.* 2018). Considering that the cataloguing of structural genomic variation is still in its infancy and that our understanding of recombination rate variation is lacking (Mérot *et al.* 2020), we adopt an inclusive approach regarding the examples discussed.

**Table 1:**
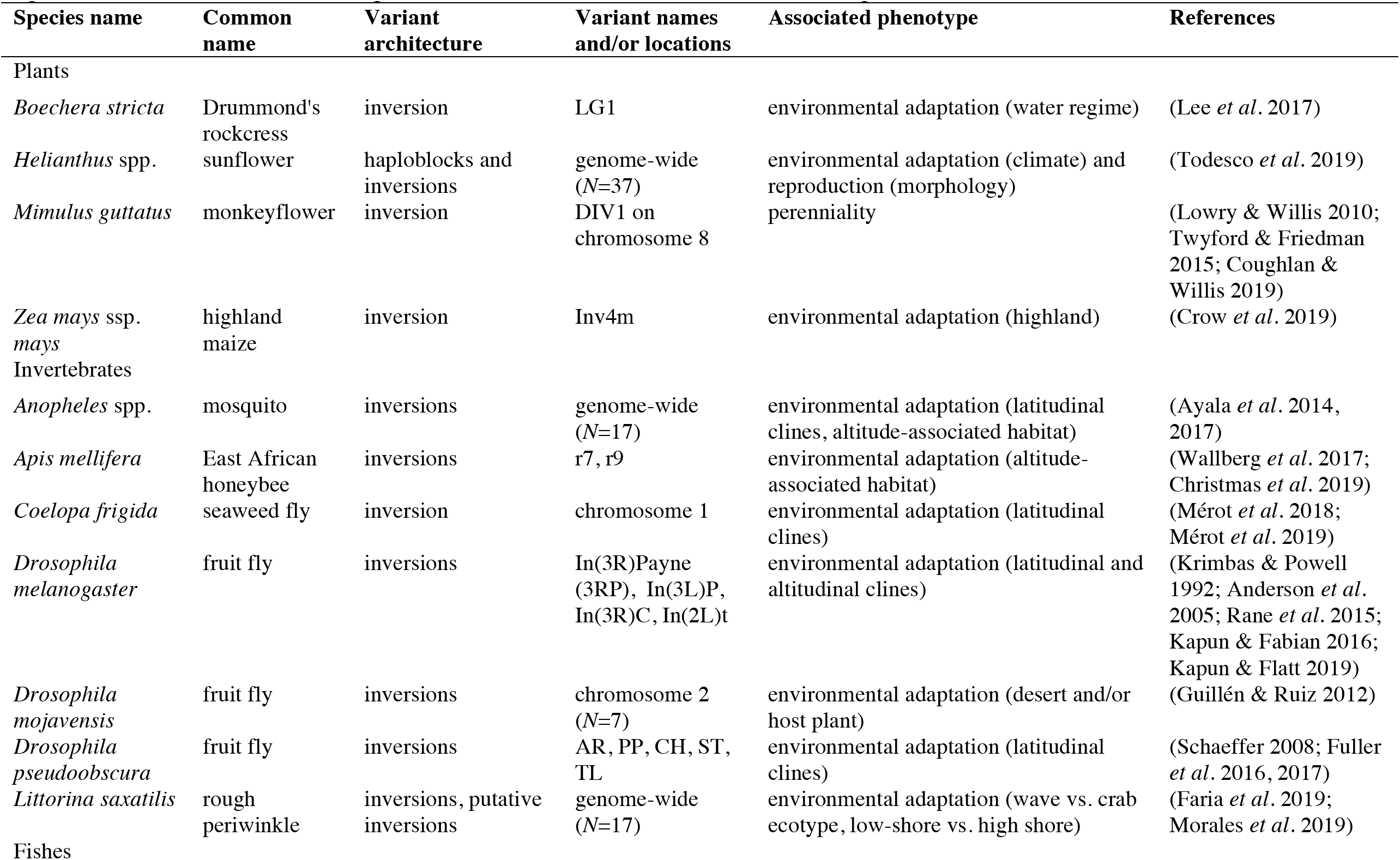
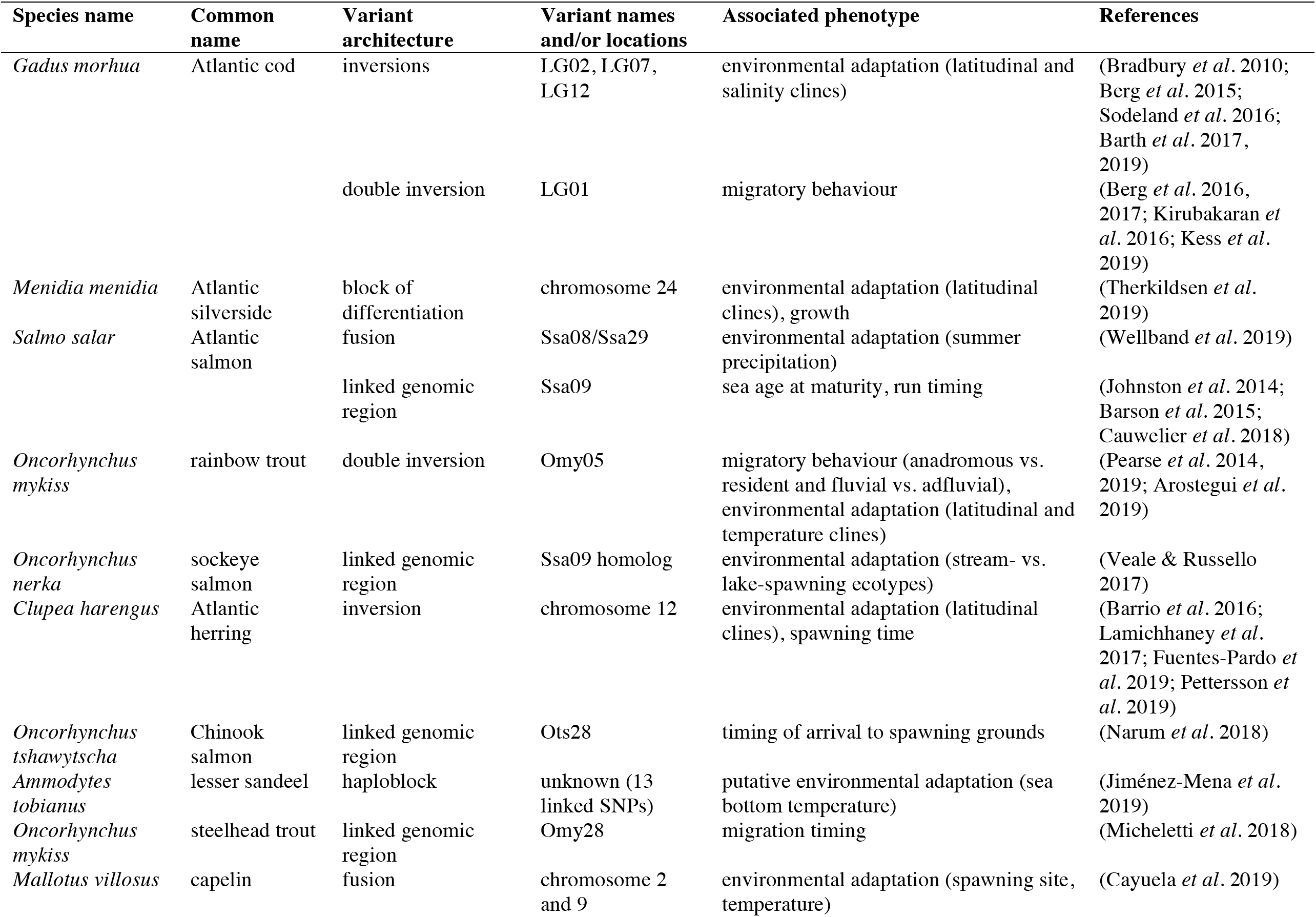
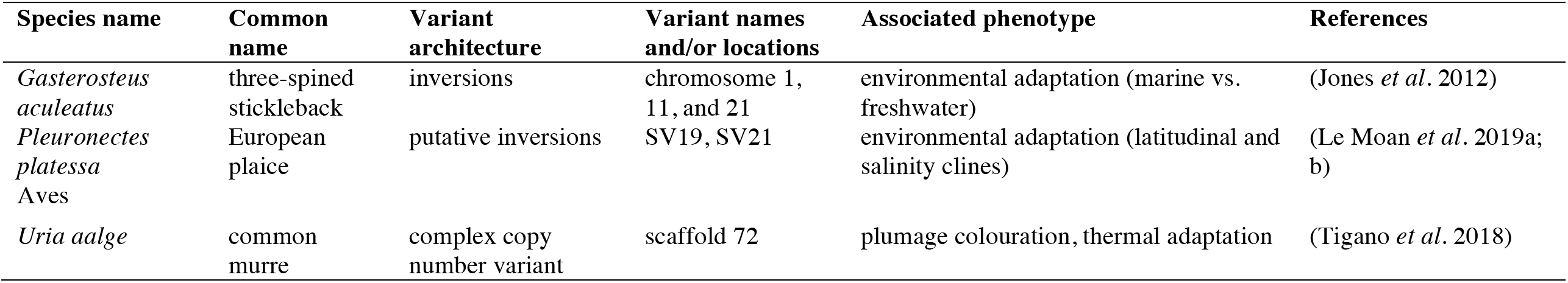
Genomic regions in linkage disequilibrium that are associated with environmental or life history traits under selection. Variant names and/or locations are specified differently across taxa and depending on the type of genomic information available. Specific names/locations are not provided if there are more than five variants per row.

### Linked architectures are favoured by selection under gene flow

Theoretical work has implicated the genetic and genomic architecture of adaptive traits as the key element in determining whether they will be lost under gene flow (Bürger & Akerman 2011; Yeaman & Whitlock 2011; Yeaman 2013; Aeschbacher & Bürger 2014; Akerman & Bürger 2014). Alleles with large effect sizes are less likely to be overwhelmed by gene flow (Yeaman & Otto 2011), as are tightly linked polygenic architectures because they effectively act as a single large-effect locus (Griswold 2006; Yeaman & Otto 2011). Gene flow increases the risk of breaking up co-adapted alleles, thereby selecting for reduced recombination and increased linkage (Nosil *et al.* 2009; Tigano & Friesen 2016).

The evolution of linked architectures under gene flow might explain why they appear to be common in species with high dispersal capabilities: flying insects, birds, and fishes (Table 1; Wellenreuther & Bernatchez 2018). Many examples of extremely tight linkage derive from systems in which closely related species or ecotypes are living in sympatry (Nosil *et al.* 2009; Hooper 2016). Yet, high gene flow is pervasive in the natural world, including many plants with high seed dispersal and marine organisms with pelagic early life stages. Therefore, linked architectures are likely common.

### Linked architectures are often directly associated with environmental adaptation

Linked architectures associated with adaptation to local environmental variables seem particularly common among flies and fishes (Table 1; Wellenreuther & Bernatchez 2018). Flies in the genus *Drosophila* provide numerous examples of inversions associated with environmental adaptation, including those exhibiting latitudinal (Krimbas & Powell 1992; Anderson *et al.* 2005; Rane *et al.* 2015; Fuller *et al.* 2016, 2017; Kapun & Fabian 2016; Kapun & Flatt 2019) and altitudinal (Kapun & Flatt 2019) clines, as well as adaptation to a desert environment (possibly through the host plant; Guillén & Ruiz 2012). Of particular interest from a human health perspective, environmentally structured inversion polymorphisms are common among *Anopheles* spp. mosquitos, potentially enhancing the adaptability and vector potency of these primary malaria vectors (Ayala *et al.* 2014, 2017). Inversions in *Anopheles* spp. vary in karyotype frequencies across latitudinal clines and between mountain forests and lowland savannahs (Ayala *et al.* 2017), and are associated with aridity tolerance.

Inversions and blocks of differentiation (which may or may not be associated with inversions) appear to underlie adaptation to salinity (Jones *et al.* 2012; Berg *et al.* 2015) and vary in frequency across latitudinal clines in marine fishes, suggesting an association with temperature or growing season length (Pettersson *et al.* 2019; Therkildsen *et al.* 2019; Kess *et al.* 2020). Chromosomal translocations and fusions are common among salmonids (Phillips 2005). Their adaptive significance, if any, is not generally known, although a fusion in Atlantic salmon (Ssa08/Ssa29) is associated with summer precipitation in a Canadian river system (Wellband *et al.* 2019).

While we have focused on adaptation to abiotic environmental variables, traits controlled by linked architectures can also be associated with the biotic environment. For example, cryptic colouration in timena stick insects (Lindtke *et al.* 2017; Lucek *et al.* 2019) and mimicry patterns in butterflies (Joron *et al.* 2013; Nishikawa *et al.* 2015), associated with a large haploblock and inversions, respectively, are under selection via local host plants and predators associated with particular environments. Changes in these environments can alter camouflage substrates and predator distributions. Adaptation will therefore depend partly on the evolutionary dynamics of the linked regions.

### Linked architectures are also indirectly associated with environmental adaptation

Architectures that link genes controlling several types of co-adapted traits can result in indirect selection on traits associated with environmental adaptation. The inversions in *Anopheles* spp. associated with differences in aridity tolerance (Cheng *et al.* 2018) are also associated with morphology and behaviour (reviewed by Ayala *et al.* 2014). Inversions underlie alternative reproductive phenotypes in some birds (Thomas *et al.* 2008; Horton *et al.* 2014; Küpper *et al.* 2015; Zinzow-Kramer *et al.* 2015), simultaneously controlling morphological (e.g., plumage colouration), behavioural (e.g., mating tactic), and life history (e.g., maturation, growth rate) traits. Common murres (*Uria aalge*) have a complex CNV maintaining linkage among genes associated with plumage colouration and thermal tolerance despite random mating (Tigano *et al.* 2018).

Alternate behavioural or life history strategies often impose different environmental selection pressures. Fish populations that migrate between freshwater and saltwater for reproduction and feeding require different temperature and salinity adaptations compared to resident populations that don’t migrate. Consequently, linked architectures in marine and freshwater fishes are associated with coexisting migratory ecotypes experiencing different environments (Table 1; Pearse *et al.* 2014, 2019; Berg *et al.* 2016; Kirubakaran *et al.* 2016; Arostegui *et al.* 2019; Kess *et al.* 2019). For example, a double inversion in steelhead/rainbow trout (*Oncorhynchus mykiss*) varies in frequency between anadromous (maturing at sea) and resident populations, as well as between fluvial (maturing in rivers) and adfluvial (maturing in lakes) populations, and exhibits latitudinal- and temperature-associated frequency clines (Pearse *et al.* 2014, 2019; Arostegui *et al.* 2019).

Variation in reproductive timing can require adaptations to different environmental conditions, such as different temperatures experienced during early life (Oomen & Hutchings 2015, 2016), which could favour architectures that link genes associated with environmental and life history traits. An inversion in Atlantic herring (*Clupea harengus*) is associated with both temperature and timing of reproduction (Fuentes-Pardo *et al.* 2019; Pettersson *et al.* 2019), whereas linked regions in Chinook salmon (*O. tshawytscha*; Narum *et al.* 2018) and steelhead (Micheletti *et al.* 2018) are associated with the timing of arrival to spawning grounds.

Many social traits in insects, birds, fishes, and plants are controlled by linked architectures due to bidirectional influences between social behaviour and genome architecture (reviewed by Rubenstein *et al.* 2019). For example, a supergene containing multiple chromosomal rearrangements underlies several social traits in the fire ant (*Solenopsis invicta*; Huang *et al.* 2018). The connection between social traits and linked architectures has broad relevance for predicting responses to environmental change, as sociality itself is often under environmental selection. For example, thermal stress and resource scarcity select for more or less sociality in different species and contexts (Doering *et al.* 2018; Kao *et al.* 2020). Further, social traits are often correlated with phenotypes that might be under selection from anthropogenic stressors such as climate change and harvesting (e.g. alternative mating tactics and growth rate in swordtail fish [*Xiphophorus* spp.; Lampert *et al.* 2010]).

Theory predicts that supergenes are also likely to arise when there is coevolution between social traits and dispersal, because dispersal will be selected against in benevolent individuals so that they tend to interact with relatives and selected for in selfish individuals so that they tend to interact with nonrelatives (Mullon *et al.* 2018; Rubenstein *et al.* 2019). Therefore, linkage between genes for dispersal traits (e.g., locomotion, physiology) and social behaviour is expected to evolve under a variety of circumstances (Rubenstein *et al.* 2019). As dispersal is one of the primary mechanisms of organismal responses to environmental change, control by linked architectures will likely alter predictions of responses to disturbance in a wide array of taxa.

Therefore, selection on diverse traits could indirectly impose selection on traits associated with environmental adaptation and incorporating linkage when modelling environmental responses has broad taxonomic utility.

## Flexible eco-genetic models can reflect a diversity of genomic architectures

The best modelling strategy will depend on what is known regarding the genomic trait architecture. For most traits, the precise architecture is not known and is often estimated to consist of between 10 and 100 unlinked loci of equal effects (e.g., Kuparinen & Hutchings 2012). Given the rising prevalence of major effect loci and tightly linked architectures, exploring both extremes – single locus, representing both major effect and tightly linked loci, and highly polygenic, unlinked loci – is warranted. In reality, mixed-architectures are likely common, and are expected to produce intermediate levels of stochasticity (Kardos & Luikart 2019). Yet, the output of these extreme scenarios will be informative about the range and distribution of possible outcomes and the sensitivity of the model to the genomic architecture in a particular case (e.g., life history or selection regime). As genomic information becomes available, more precise estimates of the number of loci, their effect sizes, and the degree of linkage among them can be incorporated into eco-evolutionary models.

## Conclusion

The challenges of obtaining estimates of LD are being rapidly overcome by high-throughput sequencing and advances in statistical genomics (Peñalba & Wolf 2020; Box 2). Our understanding of the fitness effects of linked genomic architectures is also poised for great improvements in the near future due to increased quality and accessibility of genomic resources for non-model species and, consequently, heightened interest in the topic (Mérot *et al.* 2020; Wellenreuther *et al.* 2019; Box 2). Nonetheless, simple approximations can be obtained in the meantime by treating tightly linked genomic architectures as single loci of large effect. While this approach is likely too simplistic for some purposes, it has been shown to be more powerful for detecting genotype-environment associations with linked haploblocks compared to characterizing genotypes based on SNPs within the blocks (Todesco *et al.* 2019). Ultimately, we view single-locus approximations of tightly linked architectures as a useful counterpoint to the conventional way of thinking about and modelling polygenic architectures (i.e., the quantitative genetics paradigm, which should be continually revisited in light of genomic data; Nelson *et al.* 2013). Therefore, when the genomic architecture is not known, a precautionary approach considers the greater variability and higher uncertainty that appears to be characteristic of a single-locus scenario (also see Kardos & Luikart 2019). Rapid developments in the fields of recombination rate variation and the population genomics of structural variants will continue to improve predictions borne from genomic data.

Implementing genomic architecture into spatially or temporally explicit conservation and management plans presents additional considerations. For example, variable rates of gene flow within species could result in different genomic architectures underlying adaptation at different spatial and temporal scales (Nosil *et al.* 2009; Oomen 2019). The same trait could be under selection at both scales, as contrasting genomic architectures can produce similar phenotypic outcomes (Therkildsen *et al.* 2019). The diversity of genomic architectures underlying the same or different traits also complicates the process of delineating evolutionarily significant units for conservation, particularly when the relative fitness consequences of trait variation is unclear (Waples & Lindley 2018; Waples *et al.* 2019). Nonetheless, it is clear that we must consider linked genomic architectures underlying adaptive traits when predicting the consequences of environmental disturbance to natural populations. Otherwise, by overestimating the complexity (e.g., number of independent loci) of the genomic architecture of traits under selection, we risk underestimating the complexity (e.g., nonlinearity) of their evolutionary dynamics.

## Box 1: Drift dominates evolutionary trait dynamics of single-locus architectures

We revisited the eco-evolutionary model developed by Kuparinen & Hutchings (2017) for Atlantic salmon. That model showed the evolution of age at maturity in response to fishing, assuming that it was controlled by a single locus with sexually dimorphic expression. Our objective was to compare single- and multi-locus architectures in a hypothetical species without sexually dimorphic trait expression, to illustrate how genetic architecture affects trait evolution in a more general scenario absent of sexual dimorphism. To this end, we took simple averages of the sex-specific probabilities for the male and female age at maturity in Atlantic salmon. By doing this, a male and a female carrying the same single-locus genotype have the same probabilities to mature at the ages of one sea winter (1 SW), 2 SW and 3 SW (Table I).

### Eco-evolutionary model

The model simulates the annual demographic processes of mortality (both natural and fishing), maturation, and reproduction at the level of individuals. Parameters for these processes are based on Atlantic salmon except that the probability of maturing is not sex-specific (Table I). In practice, for each individual, its survival from the previous to the next time step is determined based on binomial trials (one for natural mortality and one for fishing mortality, when applicable). For the single-locus scenario, maturation is determined based on a binomial trial using the probabilities shown in Table II. For the multi-locus scenario, maturation is based on the sum across ten loci with two alleles in each (allele sum is thus 0-20) coupled with phenotypic variability sampled from a normal distribution. Sums <6.66, 6.67-13.33, >13.34 cause maturation at the age of 1 SW, 2 SW and 3 SW, respectively. For each mature female, a mature male is sampled randomly and the number of eggs produced is based on empirical estimates (Table II). The number of eggs surviving up to grilse is sampled from binomial distributions (probabilities given in Table II). Genotypes of the newborn are sampled from the alleles of their parents and sex is assigned randomly with a 50:50 sex ratio. After these processes, the simulation proceeds to the next time step. Heritability was not held constant, but allowed to fluctuate with variation in allele frequencies. For further details about the simulation model, see Kuparinen & Hutchings (2017).

### Simulation design

To illustrate the conceptual differences between the single- and multi-locus scenarios, we simulated 10 independent evolutionary trajectories for both scenarios and tracked the average age at maturity for each simulation time step. The simulations involved three phases: i) pristine conditions in the absence of fishing (500 years, the first 400 of which were discarded as burn-in), ii) exposure to selective fishing mortality at the rate of 0.2 (selectivity given in Table II), and iii) recovery in the absence of fishing.

### Results

The simulations demonstrate that single-locus control of age at maturity generates increased variability in this trait compared to the multi-locus scenario and that the response to, and recovery from, fishing are also highly varied (Figure I). In the single locus scenario, oscillations driven by heightened genetic drift leading to chaotic dynamics are evident under pristine conditions, reduced at varying rates under fishing pressure, and markedly return in only three out of ten simulations (Figure Ia). In contrast, the multi-locus scenario does not generate chaotic dynamics and exhibits little variability both within and between simulations (Figure Ib). Consequently, single-locus control causes largely divergent and disruptive evolution of age at maturity with a wide variety of possible evolutionary trajectories and greater trait variability within trajectories, whereas polygenic control results in unidirectional evolution towards earlier maturation.

### Conclusion

Single locus control of age at maturity results in highly unpredictable evolutionary and ecological responses to fishing-induced selection on this trait, relative to the commonly assumed multi-locus control, due to greater phenotypic stochasticity (genetic drift). This appears to be the case with (Kuparinen & Hutchings 2017) and without (Figure I) sexually dimorphic expression.

## Box 2: Estimates of linkage disequilibrium and its effects on fitness are needed

### Estimates of linkage disequilibrium are becoming widely accessible

A major challenge to modelling the evolution of linked genomic architectures in diverse taxa has been the availability of relevant estimates for LD. LD depends on the distance and epistasis among loci, chromosome size, position of loci along a chromosome, local sequence content, other sources of recombination rate and gene conversion variation (e.g., genic modifiers and structural genomic polymorphisms), and demographic factors like effective population size (Figure 1; Kong *et al.* 2002; Butlin 2005; Peñalba & Wolf 2020). LD can be estimated from individual-level population-scale genomic data (Gianola *et al.* 2013; Bilton *et al.* 2018; Ragsdale & Gravel 2019). For example, population-based inference of LD can be obtained by using multiple genome alignments to calculate the statistical association of alleles, ideally accounting for demographic history (Peñalba & Wolf 2020). One advantage of this approach is that the genomic data required are already available for many systems, which also explains the frequent use of such LD estimates as proxies for inferring population recombination rates (Peñalba & Wolf 2020).

More direct estimates of recombination and gene conversion rates can be obtained using pedigree-based and gamete-based approaches (e.g., Rowan *et al.* 2019, Korunes & Noor 2019; reviewed by Peñalba & Wolf 2020). Though they can be challenging to implement in some systems (e.g., wild populations lacking pedigree information and species without external fertilization for pedigree-based and gamete-based methods, respectively), recent technological advances have made them feasible for many non-model species (Peñalba & Wolf 2020). However, biological variation in recombination rates is abundant within and among chromosomes, individuals, sexes, populations, and species, the cataloguing of which is still in its infancy (Peñalba & Wolf 2020). Importantly, even if accurate estimates of LD and/or recombination rate are obtained for a particular set of environmental conditions, recombination rates are often plastic (Stevison *et al.* 2017), complicating estimates of LD under environmental change. Nonetheless, the rapid advancement of this field will continue to eliminate the barriers associated with parameterizing LD in eco-evolutionary models.

### Linked architectures have fitness effects besides those of the target phenotype

Besides effects of the trait that is the target of selection, other aspects of linked genomic architectures impact fitness and, consequently, the evolutionary trajectory of the architecture and trait. For example, linked architectures can prevent the purging of deleterious mutations, reducing the fitness of homozygotes (Jay *et al.* 2019). Conversely, heterozygotes for linked architectures can experience partial sterility due to the production of inviable gametes during meiosis, although strong evidence for this appears limited to plants (Hoffmann & Rieseberg 2008). Whether species outside of *Diptera* can displace crossovers away from the breakpoints of chromosomal rearrangements, thus altering their patterns of inheritance, is also poorly understood, as is taxonomic variation in other types of recombination modifiers. Further, there is some probability of developing genetic incompatibilities at linked loci, which could be modelled explicitly. Finally, a better understanding of how eco-evolutionary feedbacks shape genomic architectures is needed.

## Funding

This work was supported by a James S. McDonnell Foundation 21^st^ Century Postdoctoral Fellowship Award to RAO; the Academy of Finland to AK; the European Research Council (grant number COMPLEX-FISH 770884) to AK; the Natural Sciences and Engineering Research Council of Canada Discovery Grant to JAH; the Killam Trusts to JAH; and Loblaw Companies Limited to JAH. The present study reflects only the authors’ view and the European Research Council is not responsible for any use that may be made of the information it contains.

## Acknowledgements

We thank Robin Waples for the invitation to submit this manuscript and feedback on it, as well as Marty Kardos and an anonymous reviewer for helpful comments. We also thank Marine Brieuc and Mitchell Newberry for useful discussions on aspects of the manuscript.

---

**Table I (Box 1):** Probabilities for maturation in the single-locus scenario. The values are obtained as averages from the sex-specific probabilities reported by Barson et al. (2015).

| Timing of maturity | Homozygote<br>(11) | Heterozygote<br>(10 or 01) | Homozygote<br>(00) |
| --- | --- | --- | --- |
| 2 SW → 3 SW | 0.5100 | 0.6130 | 0.9090 |
| 1 SW → 2 SW | 0.7950 | 0.2325 | 0.5660 |

**Table II (Box 1):** Demographic and fishing parameters for each age class, based on Atlantic salmon.

| Parameter | Smolt (3yr) | 1 SW | 2SW<br>(immature/mature) | 3SW+<br>(immature/mature) | References |
| --- | --- | --- | --- | --- | --- |
| survival | 0.015 (egg to smolt) | 0.07 (smolt to grilse) | 0.9/0.2 | 0.9/0.2 | Hutchings & Jones 1998;<br>Kuparinen & Hutchings 2017 |
| fishing selectivity | - | 0.15 | 0.5 | 1 | Kuparinen & Hutchings 2017 |
| eggs | - | 3040 | 7560 | 10200 | Legault 2004 |

**Figure I (Box 1):**
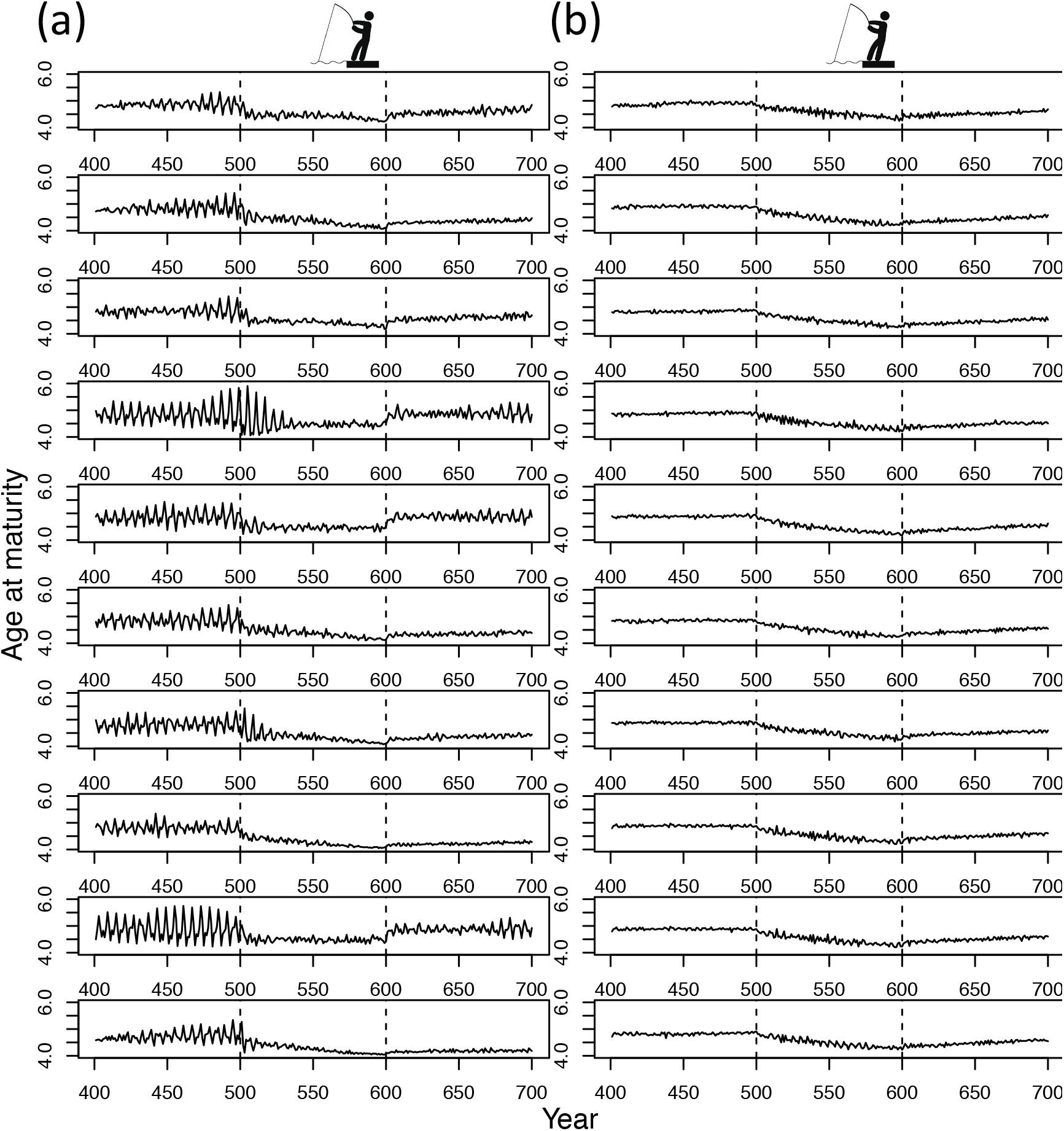
The evolution of mean age at maturity in response to fishing for a hypothetical anadromous fish population under (a) single- and (b) multi-locus scenarios for genetic architecture of the trait. Model parameters are based on Atlantic salmon except that the probability of maturing is not sex-specific for a given genotype. The beginning and end of the fishing period are indicated by dashed vertical lines. Each row represents one replicate simulation (*N*=10).

## Notes

### Competing Interest Statement

The authors have declared no competing interest.

### Summary of Updates

Version 2: A section on mixed-effect genomic architectures (combinations of major-effect and small-effect loci) has been added to address the overly simplistic dichotomy of single-locus vs. highly polygenic architectures. Importantly, the addition of small effect loci is only expected to partially alleviate the stochastic dynamics that seem to be characteristic of single locus evolutionary models. Several other minor changes have been made. Version 3: A more detailed discussion of methods for estimating linkage disequilibrium and recombination rate variation was added to Box 2, in addition to a few other minor changes.

